# Electrospun Ag-doped SnO_2_ hollow nanofibers with high antibacterial activity

**DOI:** 10.1101/629030

**Authors:** Yang Li, Xiaoning Tang, Bin Zhang, Huaming Mao

**Affiliations:** Faculty of Chemical Engineering, Kunming University of Science and Technology, Kunming 650500, Yunnan, China; Faculty of Science, Kunming University of Science and Technology, Kunming 650500, Yunnan, China

**Keywords:** Electrospinning, Ag, SnO_2_, Hollow nanofiber, Antibacterial

## Abstract

With the continuous improvement in medical science in modern times, the spread of bacterial infection has become a matter of global concern. Therefore, the search for biological medical materials with antibacterial function has become a focus of intense research. In this work, pure SnO_2_ and Ag-doped SnO_2_ hollow nanofibers were fabricated by a combination of an electrospinning method and a calcination procedure, and the effects of the doped Ag on antibacterial activity were subsequently investigated. Through the process of high-temperature calcination, a high heating rate would lead to the formation of a hollow tubular structure in SnO_2_ fibers, and Ag_2_O would be reduced to Ag0 by a facile process with appropriate thermal treatment. Additionally, the existence of SnO_2_ as a tetragonal rutile structure was confirmed. On the basis of pure SnO_2_, doping with silver greatly improved the antibacterial activity of hollow nanofibers. The formation mechanism and the antibacterial mechanism of pure SnO_2_ and Ag-doped hollow nanofibers are also discussed. This study has broad application prospects for biological medicine.

## 1. Introduction

For the foreseeable future, the threat of the drug-resistant bacteria has become one of the most serious worldwide challenges because of the growing volume of multiple novel antibiotics used in the clinic and the consequent rapid increase in antibiotic resistance [1]. During recent decades, tremendous progress has been made in antibacterial material preparation and the development of various nanofabrication techniques [2]. It has been reported that the use of nanostructured materials as novel antimicrobials has become widely accepted in the last few years and affords us a trustworthy approach for combating microbial contamination [3–5]. Antibiotics have gradually lost their bactericidal activity due to over-prescription, as bacteria have developed appropriate resistance to various antibiotics within a short period of time. As a result, inorganic nanomaterial slow releasing antibacterial agents have attracted extensive attention because of their objective advantage in terms of versatility compared with traditional antibiotics [6, 7]. Many metals, such as silver (Ag), magnesium (Mg), copper (Cu), and zinc (Zn), have historically been applied to kill bacteria long before antibiotics were widely used. Furthermore, even though a single type of metal may lose antibacterial activity, other metals can still function as antimicrobials because there is no bacterium that can develop universal resistance to all of these metals [8, 9]. In the wake of the advancement of nanotechnology, the synthesis and preparation technologies used with nanomaterials have become a research hotspot, and the physicochemical properties of metallic nanomaterials can be controlled by material scientists without inducing toxicity in human patients to produce effective antimicrobial agents [10]. The pressing trouble of antimicrobial resistance in the clinic that humans are now facing could be solved using antimicrobial metal nanoparticles designed with these considerations in mind.

Tin oxide is a good material for the degradation of organic contaminants and inhibiting bacterial growth. Tin oxide-doped nanocomposites have an increasing antibacterial capacity and photocatalytic activity in increasing concentrations [11]. Furthermore, in the preparation of composite nanomaterials, multiple preparation methods have been used, such as the absorption method, hydrothermal method, solvothermal method, and sol-gel method. In particular, the one-step method is a simple and fast synthesis method [12]. As a doping antimicrobial agent, silver has been demonstrated to have excellent antimicrobial activity and very insignificant changes in the mechanical properties of the material [13]. Previous studies have shown that silver-doped nanomaterials have extremely high bactericidal activity against *E. coli, S. aureus, E. faerial*, and *S. enterica* [11, 14].

To date, due to their unique physicochemical properties, nanofibers have been broadly researched and used in commercial applications on account of their exciting one-dimensional nanostructure. Along with the rapid progress in the synthesis and characterization of nanofibers, studies have shown that they can be used in an assortment of materials, such as polymers, carbon-based nanomaterials, and composite nanomaterials [15]. More importantly, the physicochemical properties of these metal nanomaterials can be controlled by material scientists to produce effective antimicrobial materials without exerting toxicity on human patients [10]. One-dimensional nanostructures made of tin oxide with different morphologies have been successfully prepared due to its attractive features, including low cost, nontoxicity, and ease of preparation of structures such as nanotubes [16], nanorods [17], nanobelts [18], and nanosheets [19]. Hollow tin oxide nanofibers are a kind of stable inorganic nanomaterial, and they have a unique nanometer pore structure, high surface area-to-volume ratio, and superior electrical conductivity, allowing them to supply many reaction sites and possess enhanced antibacterial activity [20]. Due to its excellent gas sensitivity, tin oxide nanofibers can be used as a sensor to detect ethanol, hydrogen, and other gases in organisms [21, 22]. The antibacterial function of these sensors is a very promising function because bacterial infection is a major problem in biological medicine. Of the various methods of creating these tin oxide nanofibers, electrospinning is a very convenient and simple preparation method for making nanofibers, which are exceptionally uniform in diameter, have large in surface area, and can be especially diversified in composition [23].

This study demonstrates the antibacterial performance of pure tin oxide hollow nanofibers and Ag-doped tin oxide hollow nanofibers created using the electrospinning method. Ag was used as a dopant in SnO_2_ hollow nanofibers, and the prepared Ag-doped hollow nanofibers were characterized in terms of both their microstructure and antibacterial properties compared to SnO_2_ hollow nanofibers using field-emission scanning electron microscope (FESEM), high-resolution transmission electron microscope (HRTEM), single point Brunauer-Emmett-Teller (BET), Energy dispersive spectroscopy (EDS), inductively coupled plasma (ICP), X-ray diffraction (XRD), and X-ray photoelectron spectroscopy (XPS). Excellent bactericidal and bacteriostatic properties were observed, which indicates the potential application of Ag-doped hollow nanofibers as high-performance antibacterial materials. In addition, the antibacterial mechanism of the above hollow nanofibers is also discussed.

## 2. Materials and Methods

### 2.1 Materials

Polyvinylpyrrolidone (PVP, Mw = 1,300,000), ethanol (purity > 99.7 wt%), stannic chloride dehydrate (SnCl_2_·2H_2_O), silver oxide (Ag_2_O), N,N-dimethylformamide (DMF), and sodium chloride (NaCl) were purchased from Tianjin Fengchuan Chemical Reagent Technology Co. Ltd., and all of the above chemicals were of analytical reagent grade (AR). The beef paste (Total nitrogen > 13 wt%), peptone (Total nitrogen > 14.5 wt%), and agar powder (Solid sexual content > 77%) used in the agar culture medium were all from Beijing Aobox Biotechnology Co. Ltd. Select gram-negative bacteria (*E. coli*) were purchased from China Center for Type Culture Collection (CCTCC, AB 204033), and gram-positive bacteria (*S. aureus*) were purchased from School of Life Science, Kunming University of Science and Technology.

### 2.2 Preparation

Two different materials, pure SnO_2_ and Ag-doped hollow nanofibers, were synthesized using electrospinning technology. A total of 0.700 g SnCl_2_·2H_2_O and 0.002 g Ag_2_O were added to DMF to obtained reagent A, and 0.700 g PVP was dissolved in 5 mL ethanol to obtain regent B. Afterward, A was stirred rapidly for 1 h at 35°C, and then both A and B mixed together by stirring at constant speed for several hours at 35°C, resulting in the prepared precursor solution. The precursor solution was then loaded into a glass syringe for electrospinning at 13.3 kV, after which, the collected products were put into a tube furnace, and the organic ingredients were removed by heating at high temperature, resulting in the Ag-doped hollow nanofibers. The decomposition conditions of Ag_2_O are given. In addition to the addition of elemental Ag, pure SnO_2_ hollow nanofibers were synthesized by the same method.

### 2.3 Materials characterization

X-ray diffraction (XRD, D/Max-3B), which was measured with Cu Kα_1_ radiation (λ = 1.5406), scanning over 10–90°, was used to investigate the crystalline structure of the samples. The surface morphologies of pure SnO_2_ and Ag-doped hollow nanofibers were investigated using a field emission scanning electron microscopy (FESEM, SU-8010) and transmission electron microscopy (TEM). Energy-dispersive spectroscopy (EDS) was used in conjunction with SEM at an acceleration voltage of 10 kV. The valence state of the Ag ions in the composites was analyzed using X-ray photoelectron spectroscopy (XPS, ESCALAB 250-Xi). The Brunauer-Emmett-Teller (BET) specific surface areas of the samples were revealed on a JW-BK222 specific surface area and aperture analyzer.

### 2.4 Antimicrobial performance

The antimicrobial performance of pure SnO_2_ and Ag-doped hollow nanofibers was quantitatively analyzed using *E. coli* and *S. aureus* in terms of bacterial growth curves and minimum inhibitory concentration (MIC) following the standard antibacterial activity protocol (GB/T 21510-2008, oscillation method) for nano-inorganic antibacterial materials [24]. The lowest concentration of the Ag-doped hollow nanofibers that inhibited the bacterial growth completely was known as the most appropriate definition for the MIC. All tests were carried out in triplicate.

The initial bacterial count was kept between 5 and 6 ×10^6^ CFU/mL after dilution, and afterward, samples were plated on agar culture media and incubated for 18–22 h at 37°C. All bacterial data were taken from agar culture media using a bacterial colony counter. The antibacterial rate was defined as

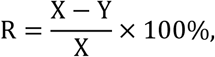

where R is the antibacterial rate, X is the number of colonies in control sample, and Y is the number of bacterial colonies in the test sample.

## 3. Results and Discussion

### 3.1 Structural and morphological analysis

XRD was performed, and all the peaks for pure SnO_2_ and Ag-doped hollow nanofibers are shown in Fig. 1(a). After comparing these with the standard spectrogram JCPDS card no. 41-1445, the crystal structure of both was indexed as a single-phase tetragonal rutile SnO_2_ structure. The effect of doping on the crystallinity of SnO_2_ hollow nanofibers was obtained by monitoring the diffraction peaks. The spectra displayed that there was a handful of changes in the (200) peaks and the (112) and (321) diffraction peaks and a slight left shift in the (110) and (101) diffraction peaks of Ag-doped hollow nanofibers as compared with those of SnO_2_ hollow nanofibers. This possibly suggests that the incorporation of Ag results in a lattice deformation of SnO_2_. The crystallite size of pure SnO_2_ and Ag-doped hollow nanofibers was estimated to be 28.m and 37.9 nm, respectively, according to the Debye-Scherrer equation (*D* = 0.89*γ*/*β*cos*θ*)[25]. In fact, this change in the XRD diffraction peaks of the Ag-doped hollow nanofibers can be attributed not only to Ag incorporation but also to changes in crystallite size. Thus, in this study, it was not clear what caused the shift in diffraction peak position or whether Ag has replaced Sn in the SnO_2_ lattice [26]. Notably, an extremely high purity of the pure SnO_2_ and Ag-doped hollow nanofibers without any byproducts occurred because there were no other diffraction peaks as shown in this figure. However, no characteristic diffraction peaks of Ag_2_O or Ag^0^ were detected, possibly due to the low content of Ag_2_O or Ag^0^ [27]. Fig. 1(b) shows a simulated primitive cell of the SnO_2_ structure. In SnO_2_, each oxygen ion is bound to three tin ions and is surrounded by six oxygen ion atoms. Ag may replace Sn in the crystal cell and change the lattice parameters, which can be calculated [28].

**Fig. 1.**
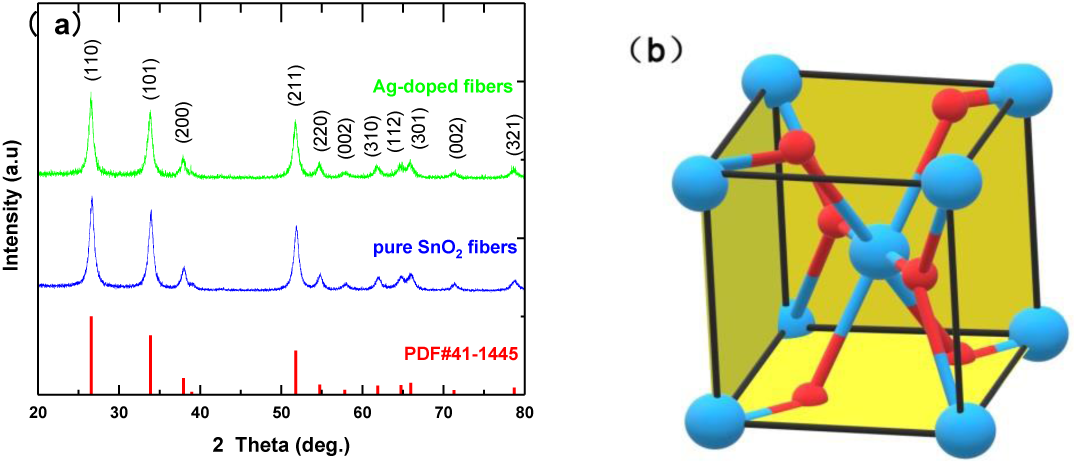
(a) X-ray diffraction patterns of non-doped and Ag-doped hollow nanofibers. (b) Unit cell of SnO_2_ (Blue is Sn atoms, and red is O atoms).

Figures 2(a) and (b) show SEM images of the SnO_2_ hollow nanofibers and the Ag-doped hollow nanofibers after removal of organic components. These nanofibers are rod-like structures composed of a collection of uniform tiny particles, and these particles were about 39–76 nm in diameter. The diameter of the nanofibers was about 119–223 nm, and the length reached the millimeter level, indicating a large aspect ratio. The illustrations in (a) and (b) are high power SEM images, displaying the differences between the two samples. The high power images clearly show the unique hollow structure of the nanofibers. There was a rougher surface on the Ag-doped nanofibers compared to the pure SnO_2_ nanofibers. In other words, the doping of Ag resulted in the rougher surface of the nanofibers, which may be attributed to the fact that Ag replaced Sn in the SnO_2_ lattice. Table 1 shows the fiber diameter, grain size, specific surface area, and bore diameter of pure SnO_2_ nanofibers and Ag-doped nanofibers. The addition of Ag reduces the diameter of the fiber and the diameter of the particles that make up the fiber [29, 30]. This also was confirmed by BET measurements of changes in the microstructure caused by the Ag addition. The specific surface area of Ag-doped nanofibers increased significantly because of the decrease in grain size. The increase of specific surface area can effectively increase the amount of Ag^+^ released by nano-Ag, so the antibacterial effect was significantly enhanced. However, the change in aperture size cannot be explained by the addition of Ag and may be an error in measurement. The EDS images [Fig. 2(c)] show the approximate amounts of various elements. Five characteristic peaks appeared (C, O, Ag, Sn, and Au), and the other unlabeled peaks were assigned to Si and N, indicating that nano-Ag was successfully doped into the SnO_2_ nanofibers. The presence of Au and C is the basic requirement for conducting electricity. The content of Ag was about 1% that of Sn, which was related to the amount of reagent added during the preparation process (see Materials and Methods).

**Table 1.**
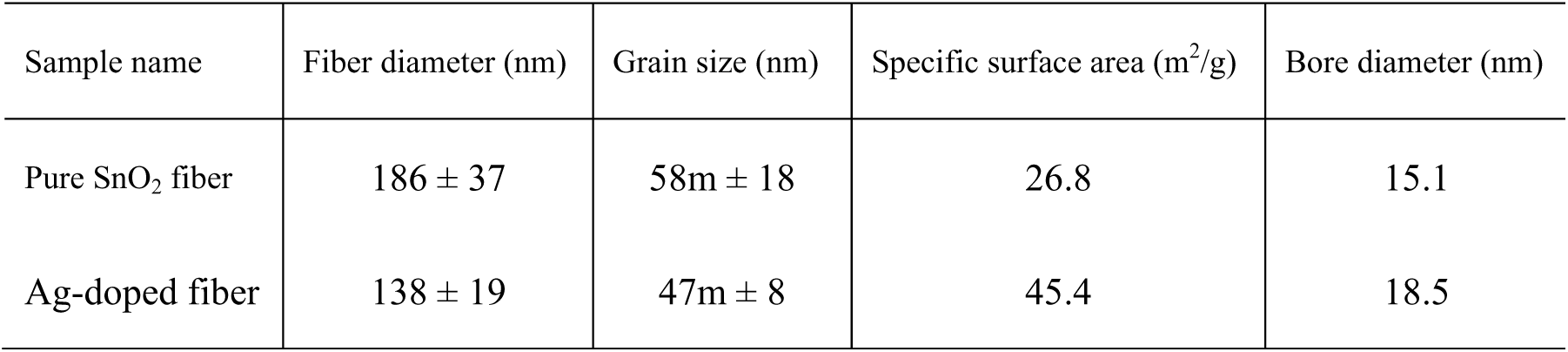
Fiber diameter, grain size, specific surface area, and bore diameter of pure SnO_2_ nanofibers and Ag-doped nanofibers.

**Fig. 2.**
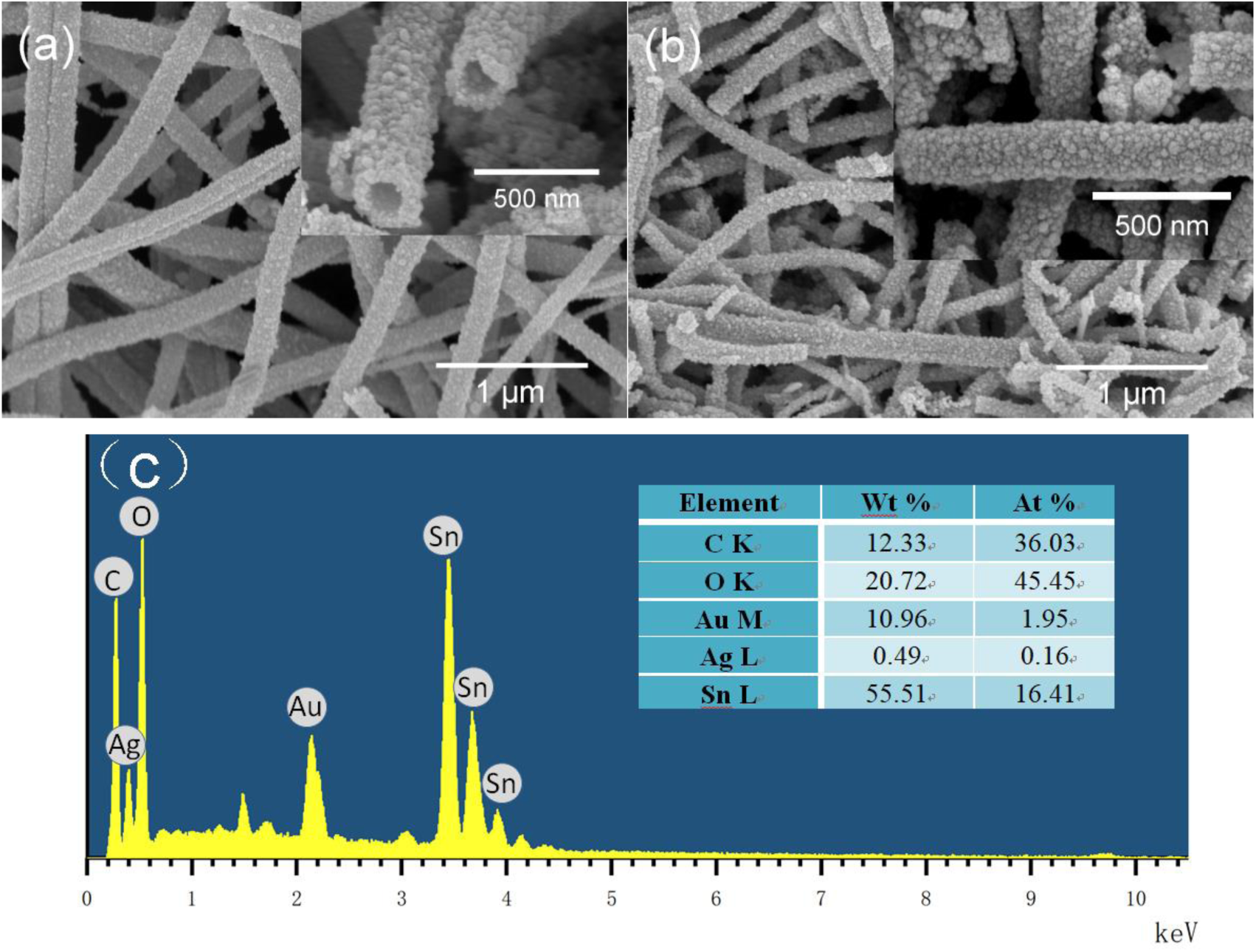
SEM images of (a) pure SnO_2_ nanofibers and (b) Ag-doped SnO_2_ nanofibers. (c) EDS images of Ag-doped nanofibers. The insets show corresponding enlargements of nanofiber sections.

TEM and HRTEM were used to characterize the nanofibers in order to investigate the microstructure of the SnO_2_ samples in greater detail. Fig. 3(a), (c), and (e) shows the TEM and HRTEM results for pure SnO_2_ nanofibers, and Fig. 3(b), (d), and (f) shows Ag-doped nanofibers. The hollow nanofiber structure was consistent with that seen in the SEM images. The nanofibers from Ag-doped nanofibers had a rougher and denser surface and smaller particles compared with pure SnO_2_ nanofibers. High-resolution transmission electron microscopy (HRTEM) was used to further verify the grain sizes of pure SnO_2_ and Ag-doped nanofibers. The average particle size of a pure SnO_2_ fiber and an Ag-doped fiber was 57 nm and 45 nm, respectively. The results indicated that the grain size to decrease is due to the addition of Ag. The HRTEM images (Fig. 3(e) and (f)) indicate the pure SnO_2_ and Ag-doped hollow nanofibers are polycrystalline structures because the lattice distance of 0.33 nm, 0.318 nm and 0.263 nm, respectively, shown in the images, corresponded to the (110), (001), and (101) lanes of the rutile (tetragonal) SnO_2_ structure [31]. In addition, some of the small particles in the figure may be nano-Ag.

**Fig. 3.**
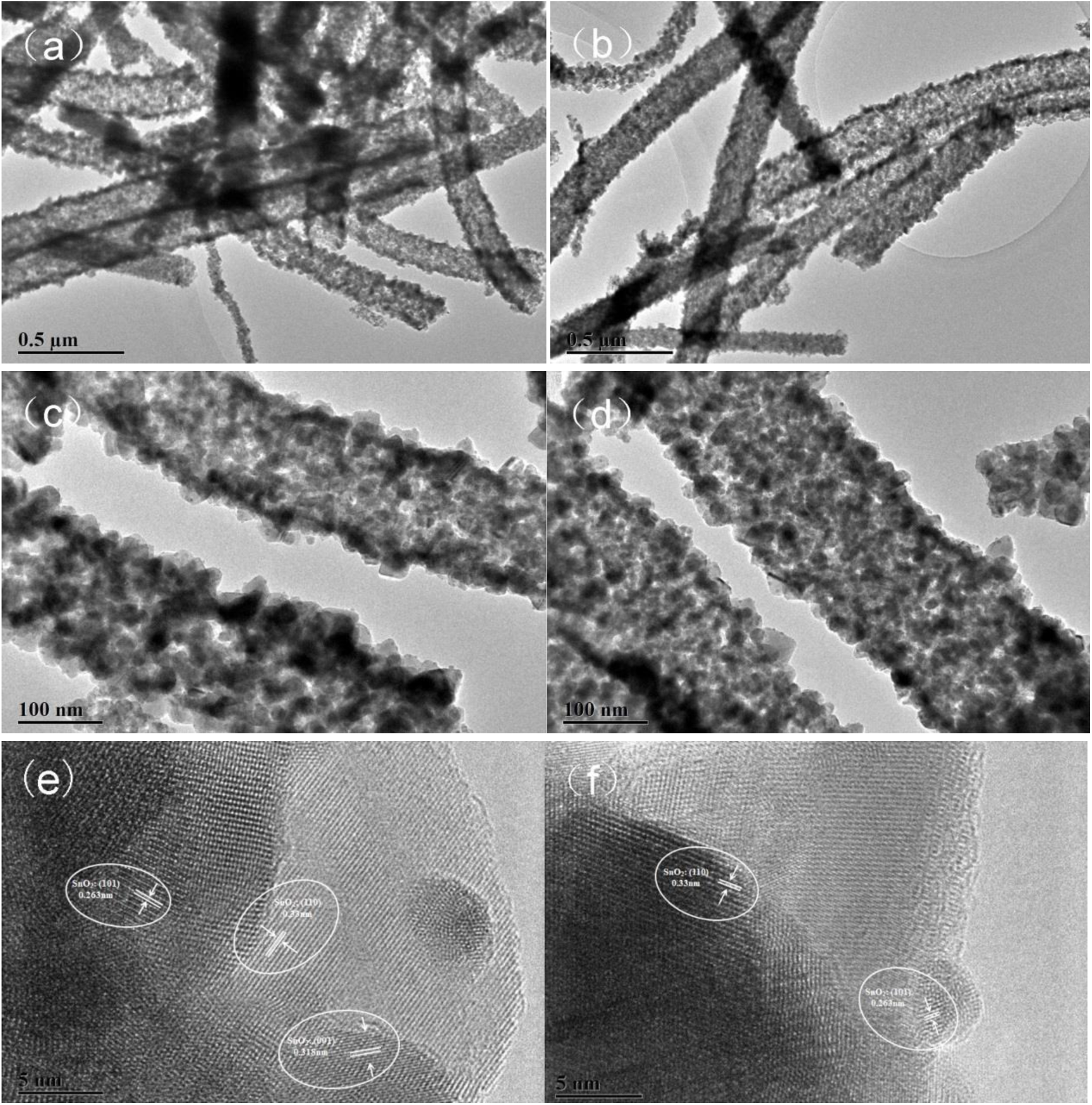
(a and c) TEM images and (e) HRTEM image of pure SnO_2_ nanofibers, and (b and d) TEM images and (f) HRTEM images of Ag-doped nanofibers.

### 3.2 XPS analysis

In order to determine the surface compositions and chemical state of the as-prepared Ag-doped SnO_2_ hollow nanofibers, XPS spectra of the materials were determined and are shown in Fig. 4, and nothing else was observed in the spectra other than the composition of the materials. These results were similar to the EDS analysis results, and the C 1s peak was from the spectrometer instrument. The XPS spectrum of the Ag-doped hollow nanofibers is shown in Fig. 4(a). The central binding energies of O 1s, Sn 3d_5/2_, and Sn 3d_3/2_ peaked at 530.6 eV, 486.5 eV, and 495.0eV, respectively, which was very close to the value reported in the standard spectrum of the lattice of oxygen and tin in SnO_2_ [32]. Moreover, the spin-orbital coupling energy gap between the Sn 3d_5/2_ and Sn 3d_3/2_ energy levels was about 8.5 eV, which is in good agreement with the values reported in the standard spectrum of Sn 3d. The 3d spectrum of Ag 3d contained two symmetric peaks at 368.3 eV and 374.3 eV. When the spin separation was 6.0 eV, the two peaks belonged to Ag 3d_5/2_ and 3d_3/2_ binding energies [33]. These results show that after calcining at high temperature, Ag was reduced to Ag^0^ and existed in the SnO_2_ nanofibers. Combining EDS [Fig. 2(c)] and XPS [Fig. 4(d)], it can be seen that a small amount of AgO and Ag_2_O existed, which meant that not all Ag_2_O was reduced to Ag^0^ during nanofiber preparation [34].

**Fig. 4.**
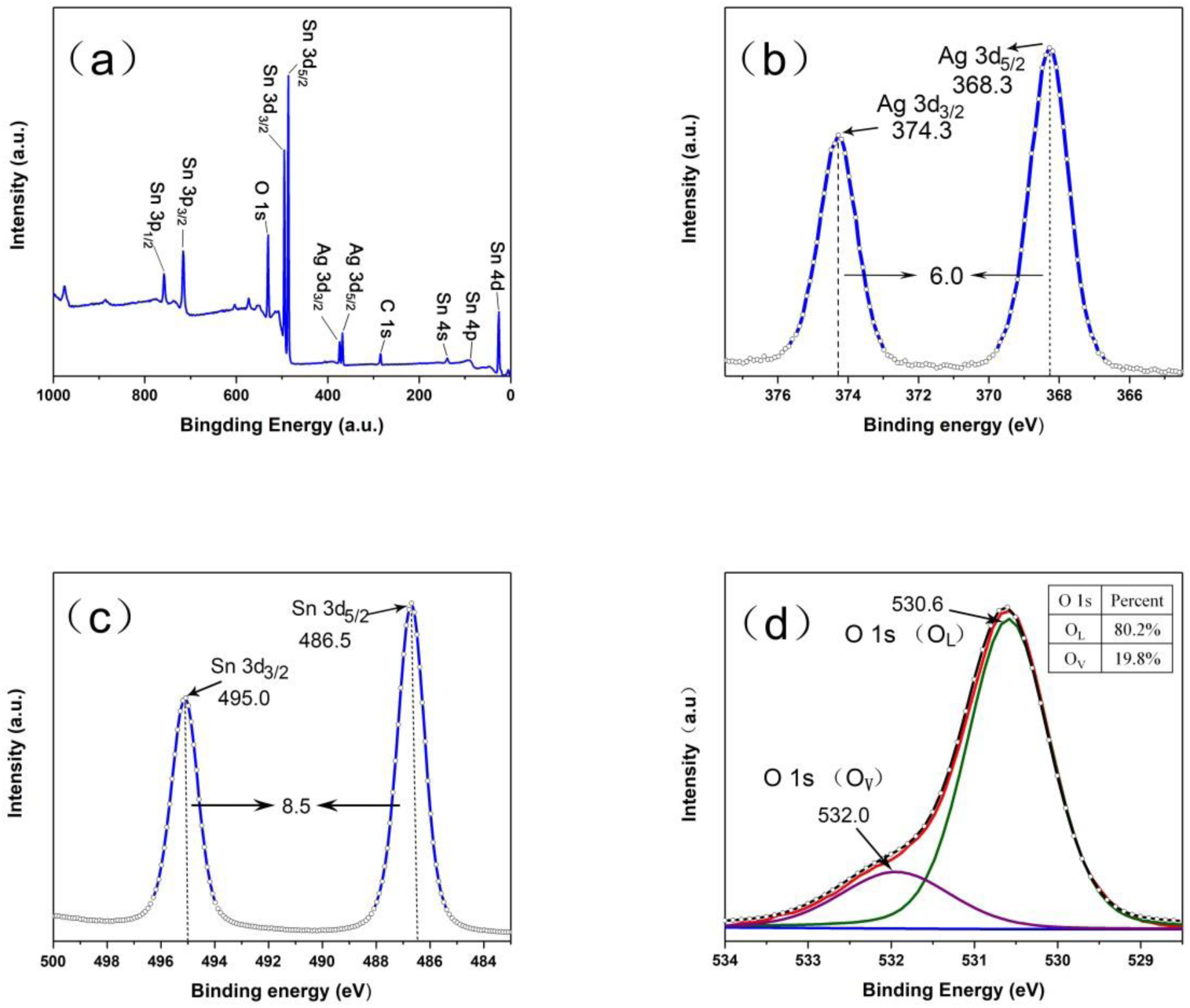
XPS spectra of Ag-doped SnO_2_ hollow nanofibers, (a) survey spectrum and high-resolution spectra for (b) Sn 3d, (c) Ag 3d, and (d) O 1s.

### 3.3 Antimicrobial performance analysis

A cell viability test was based on using different concentrations of pure SnO_2_ and Ag-doped hollow nanofibers [Fig. 5(a)]. As shown in Fig. 5(a), there was a variable performance of different materials on cell survival. These experiments each used 10 mL of the same concentration of cell solution. The concentration of the cell solution of *E. coli* and *S. aureus* was 5.5 ± 0.5×10^6^ CFU/mL and 5 ± 0.4×10^6^ CFU/mL, respectively. Pure SnO_2_ have poor antibacterial activity, not only toward *E. coli* but also toward *S. aureus*, and with increasing concentration of SnO_2_ hollow nanofibers, there was only a slight increase in the antibacterial rate. By comparison, the antibacterial effect of the Ag NP-containing SnO_2_ nanomaterials improved dramatically at each concentration. The antibacterial rate with Ag-doped nanofibers was as high as about 98% when 100 mg of material was added, which was about three times of that of materials without Ag-doping at the same concentration of the bacteria solution of *E. coli*. The antibacterial effect of the two kinds of materials was less due to the special structure of *S. aureus* compared to *E. coli*, but the Ag-containing materials still had great advantages relative to pure SnO_2_ nanofibers.

**Fig. 5.**
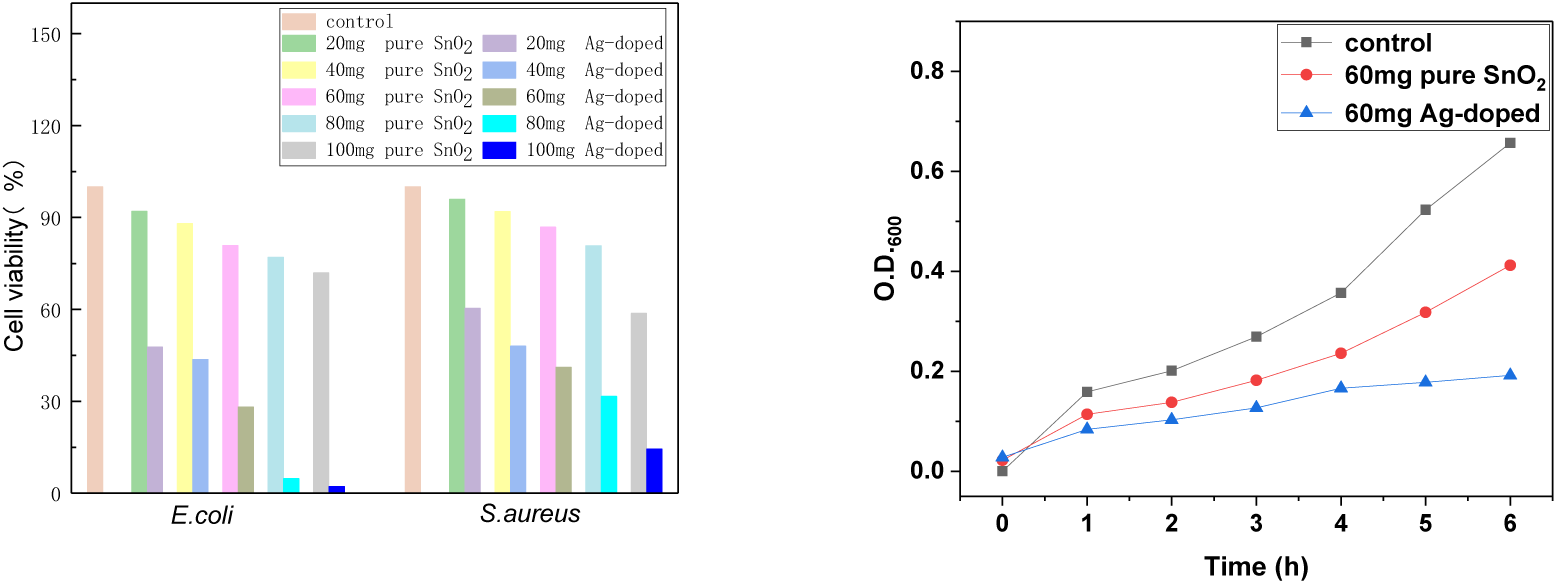
(a) Cell viability of *E. coli* (left) and (b) *S. aureus* (right) treated with control or different concentrations SnO_2_ and Ag-doped hollow nanofibers. (b) Cell growth of *E. coli* on the control, 60 mg pure SnO_2_, or 60 mg Ag-doped hollow nanofibers over time.

The absorbance of bacterial solution with time was measured by a UV spectrophotometer [Fig. 5(b)]. The *E. coli* cell reached exponential growth after 6 h of incubation in the case of the experimental control. In addition, the cell solution with added SnO_2_ hollow nanofibers also showed rapid growth after 6 h of incubation. In the beginning, the bacteria in the Ag-doped hollow nanofibers added solution showed an obvious upward trend; however, the growth of the bacteria was gradually inhibited, and the change curve in the OD value tended to be flat over time. The slight increase in the initial absorbance may have been related to the addition of the SnO_2_ materials. Photographs of bacterial culture plates are shown in Fig. 6, allowing visualization of the survival rate after 0, 1, 2, 3, 4, 5, and 6 h of exposure to the pure SnO_2_ or Ag-doped nanofibers. The growth trend was consistent with that shown in Fig. 5(b). *E. coli* and *S. aureus* cells used above were allowed to grow at 37°C to early exponential phase.

**Fig. 6.**
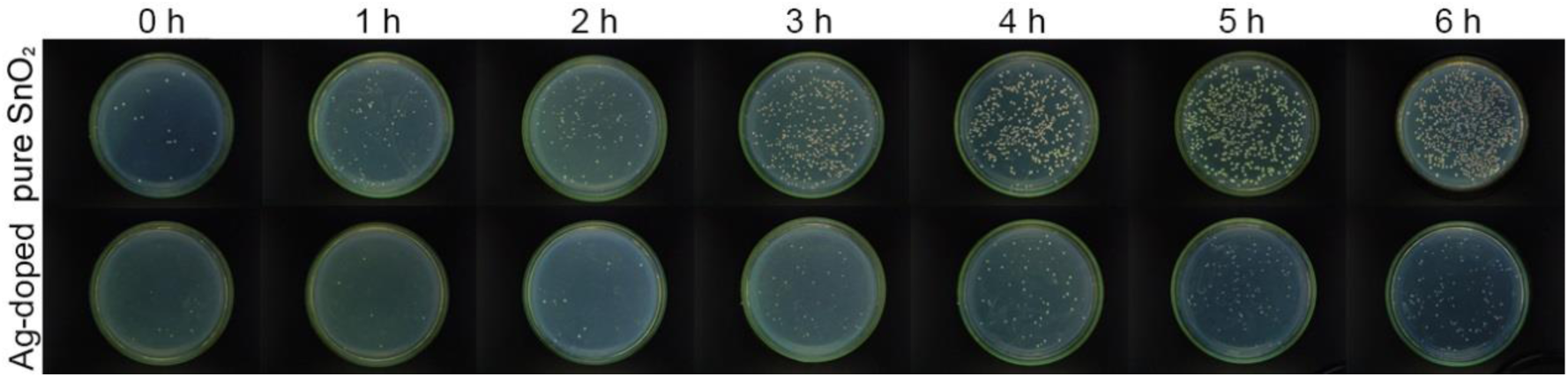
*E. coli* growth in the presence of Ag-doped or pure SnO_2_ nanofibers.

### 3.4 The possible formation mechanism of the hollow nanofibers

Previous studies have shown that the formation of the hollow nanofibers was governed by solvent evaporation rates and phase separation [35]. The faster the solvent evaporates, the more obvious the hollow nanofiber skin layer [36]. This is shown as the possible formation mechanism of the hollow nanofibers in Fig. 7. The PVP and SnCl_2_ were dissolved in a solvent consisting of DMF and distilled water, and the elemental Sn was ionized to Sn^4+^ in the water. Ethanol not only played the role of solvent while particles were dissolved but also played a role in promoting phase separation because of its volatility. Some smaller particles, such as Sn^4+^ and Ag_2_O, gradually moved toward the boundary layer, while larger PVP molecules were wrapped by small molecules to form the double-layer tubular structure as shown in Fig. 7. Ethanol evaporated continuously at the fiber boundary, and phase separation occurred during this process. Finally, hollow nanofibers were formed by the calcining and decomposition of PVP and Ag_2_O broken down into Ag^0^.

**Fig. 7.**
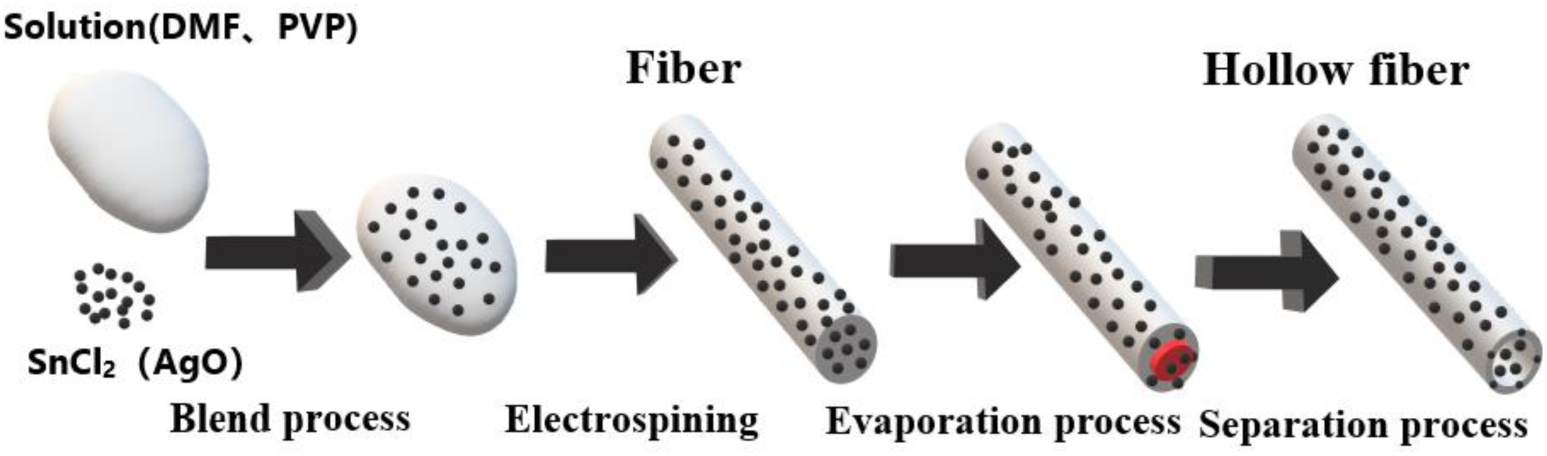
Schematic of the formation mechanism of hollow nanofibers.

### 3.5 The possible antibacterial mechanism of Ag-doped hollow nanofibers

The agglomeration effect of Ag NPs had a significant effect on antibacterial activity, and a good dispersion of nanoparticles was required for effective antibacterial activities [37]. Nano-level antimicrobial materials have excellent antimicrobial activity, but these are usually used in combination with some measures to prevent their tendencies to agglomerate [38], such as adding Bovine serum albumin [39] or β-cyclodextrin [40]. As nanofibers are one-dimensional nanomaterials, the agglomeration effect is not significant. So the antibacterial material prepared by doping Ag in nanofibers can stably release Ag^+^ at the nanolevel, thus achieving excellent antibacterial effects.

Four antibacterial processes in *E. coli* cell are outlined in Fig. 8. Smaller Ag particles help release Ag^+^ more quickly. The size of the Ag particles directly affects the effective surface of Ag and then affects the dissolution rate of Ag and the resulting antibacterial activity [41]. The separation of the membrane from the cytoplasm of Ag-treated *E. coli* may be the cause of cell death. The substantial negative charge on the cell membrane is neutralized by positively charged Ag^+^, and the destruction of the cell membrane results in the leakage of the cytoplasm [42]. When Ag reacts with the cell membrane, it is easy to release Ag^+^, which then enters the cell and may damage DNA, causing mutation or death [43]. It has also been reported that Ag NPs can destroy cell mitochondria and cause cell death due to insufficient energy supply. Another cause of cell death is the destruction of the internal structure of cells by the interaction of Ag^+^ with the ROS produced by cells [44].

**Fig. 8.**
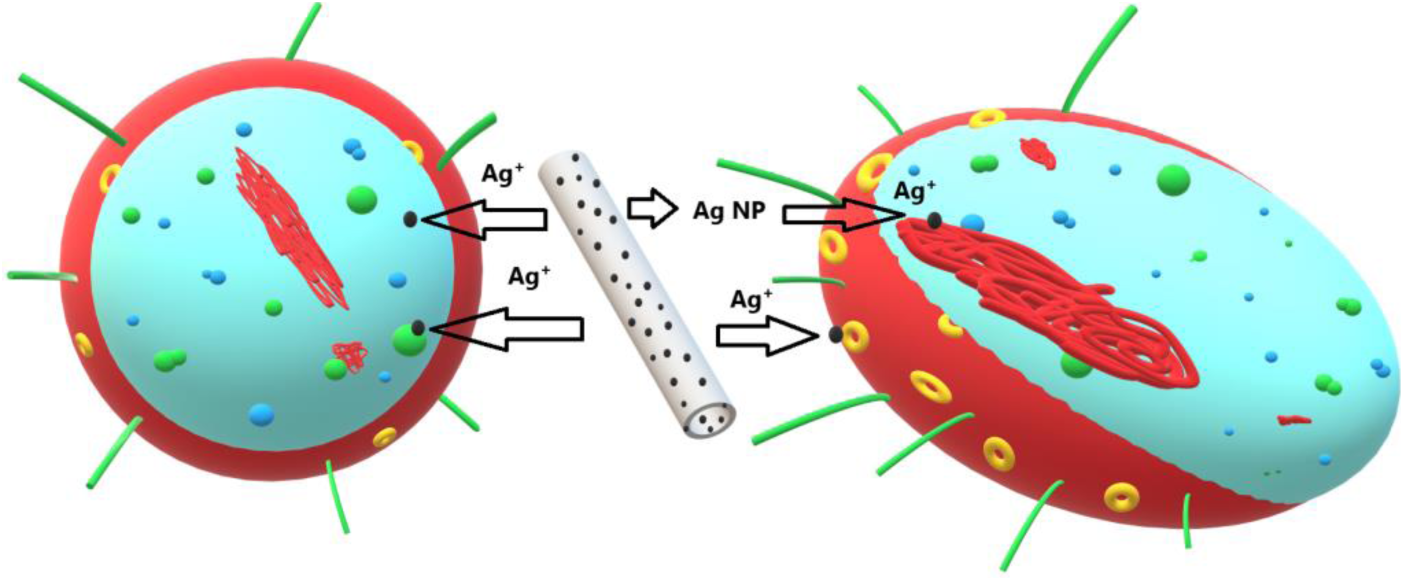
Schematic of the antibacterial mechanism of Ag-doped hollow nanofibers.

## 4. Conclusions

In summary, tubular Ag-doped SnO_2_ hollow nanofibers were prepared by electrostatic spinning and after calcination. No contamination with other impurities was detected in the prepared materials, but a small amount of Ag_2_O did not change into Ag0. We also confirmed that the SnO_2_ existed in a tetragonal rutile structure. Using both gram-negative (*E. coli*) and gram-positive (*S. aureus*) bacteria to test antimicrobial activity, we found that the Ag-doped nanofiber antimicrobial activity was far higher than that of pure SnO_2_ nanofibers, and the antibacterial rate versus *E. coli* antibacterial was more than 99% for 100 mg (5 mg/mL), while for *S. aureus*, the antibacterial rate was above 80%. It was worth mentioning that Ag-doped hollow nanofibers had a smaller grain size, less agglomerative effects, larger specific surface area to release bactericidal Ag ion, and near perfect bactericidal effects, meaning that Ag-doped hollow nanofibers have good prospects for use in the field of antibacterial and biomedical materials.

## Acknowledgments

This work was supported by a grant from the National Natural Science Foundation of China, China (21677010). We thank LetPub (www.letpub.com) for its linguistic assistance during the preparation of this manuscript.

